# Auditory detection is modulated by theta phase of silent lip movements

**DOI:** 10.1101/2020.07.07.186452

**Authors:** Emmanuel Biau, Danying Wang, Hyojin Park, Ole Jensen, Simon Hanslmayr

## Abstract

Audiovisual speech perception relies, among other things, on our expertise to map a speaker’s lip movements with speech sounds. This multimodal matching is facilitated by salient syllable features that align lip movements and acoustic envelope signals in the 4 - 8 Hz theta band. Although non-exclusive, the predominance of theta rhythms in speech processing has been firmly established by studies showing that neural oscillations track the acoustic envelope in the primary auditory cortex. Equivalently, theta oscillations in the visual cortex entrain to lip movements, and the auditory cortex is recruited during silent speech perception. These findings suggest that neuronal theta oscillations may play a functional role in organising information flow across visual and auditory sensory areas. We presented silent speech movies while participants performed a pure tone detection task to test whether entrainment to lip movements directs the auditory system and drives behavioural outcomes. We showed that auditory detection varied depending on the ongoing theta phase conveyed by lip movements in the movies. In a complementary experiment presenting the same movies while recording participants’ electro-encephalogram (EEG), we found that silent lip movements entrained neural oscillations in the visual and auditory cortices with the visual phase leading the auditory phase. These results support the idea that the visual cortex entrained by lip movements filtered the sensitivity of the auditory cortex via theta phase synchronisation.

## INTRODUCTION

When hearing gets difficult, people often visually focus on their interlocutors’ mouth to match lip movements with sounds and to improve speech perception. Mouth opening indeed shares common features with auditory speech envelope, which temporally synchronize on dominant 4 - 8 Hz theta rhythms imposed by syllables (Park et al., 2016; Chandrasekaran et al., 2009; Luo & Poeppel, 2007). The present study focuses on theta activity conveyed by moving lips because the speaker’s mouth provides a direct source of visual speech information matching sounds. Nevertheless, we acknowledge that other visual cues like the eyebrows convey features that structure the speech signal as well but at different dominant time-scales (e.g., 0.5-3Hz delta prosodic features; Ghitza, 2017). In the brain, neural oscillations from the auditory cortex track the auditory envelope structure during speech perception, suggesting that this “entrainment” reflects signal analysis (Keitel et al., 2018; Pelle & Davis, 2012; Gross et al., 2013; Giraud & Poeppel, 2012). Although the term entrainment is currently under debate (Meyer, Sun & Martin, 2019; Obleser & Keyser, 2019; Haegens & Zion Golumbic, 2018; Rimmele et al., 2018), here we use it to describe neural patterns tracking salient features conveyed in speech signals which occur at theta frequency (4-8 Hz).

Previous studies demonstrated that the perception of moving lips entrains oscillations in the visual cortex and modulates activity in the auditory regions, although not limited to the theta band (Bourguignon et al., 2020; Crosse et al., 2019). Further, information specific to lip movements is represented not only in the visual cortex but also in the auditory cortex (Park et al., 2018). In an fMRI study, Calvert et al. (1997) reported overlapping activations in the bilateral primary auditory cortex during the perception of isolated words presented either visually (i.e., silent moving lips) or aurally. More recently, Besle et al. (2008) used intracranial recordings in epileptic patients to investigate the neural responses evoked by the perception of lip movements in the auditory cortex during the presentation of syllables in uni- or multimodal conditions. They reported activations in response to silent lip movements in the visual cortex followed by similar responses in the secondary auditory cortex, suggesting crossmodal activation via direct feedforward processes. However, all these results beg the question of whether visual perception of lip movements modulates the auditory cortex in a functional way. In other words, do purely visually induced theta speech rhythms impose time windows that render the auditory cortex more sensitive to inputs in a phasic manner? If the answer to this question is yes, then visually focussing on your interlocutor’s mouth when you have trouble understanding them would indeed be an effective filter modulator to increase auditory sensitivity.

## RESULTS

### Entrainment to lip movements during silent speech drives behavioural performance

To address this question, we adapted an auditory tone detection paradigm in which a continuous white noise was presented simultaneously with silent movies displaying speakers engaged in conversations (Figure 1 and see Material and Methods; Movie1 and Sound1 for examples). Participants were instructed to press a key as fast and accurate as possible every time when they detected a pure tone (1 kHz, 100 ms) embedded in the white noise at individual threshold (determined with a calibration task). In the condition of interest, there were two target tones: the first tone occurred randomly in the first half of the trial (0 to 2.5 s after trial onset) and the second tone occurred randomly in the second half of the trial (2.5 to 5 s). Two additional conditions containing zero or one single tone were introduced to estimate the false alarm rates (FA) and to reduce the predictability of the second tone by the occurrence of the first one. The three conditions were counterbalanced and randomised across six blocks of 50 trials (100 trials per condition). To test the first hypothesis of visual entrainment affecting auditory processing, participants were asked to attend carefully to the silent movies centred on the speakers’ nose displayed with sound albeit non-informative. Crucially, the videos were preselected such that lip movements occurred in the 4 - 8 Hz theta range. We determined at which theta frequency the vertical mouth’s apertures and auditory speech envelope showed significant dependencies in the original clips by using mutual information method (see Material and Methods section). This paradigm allowed us to link directly the onset of detected tones with the phase of the ongoing theta activity conveyed by the lip movements. As neural entrainment increases over time (Thut et al., 2011; Hanslmayr, Axmacher & Inman, 2019), we compared behavioural performance between the early and late time-windows (containing respectively the first and second tones).

**Figure 1:**
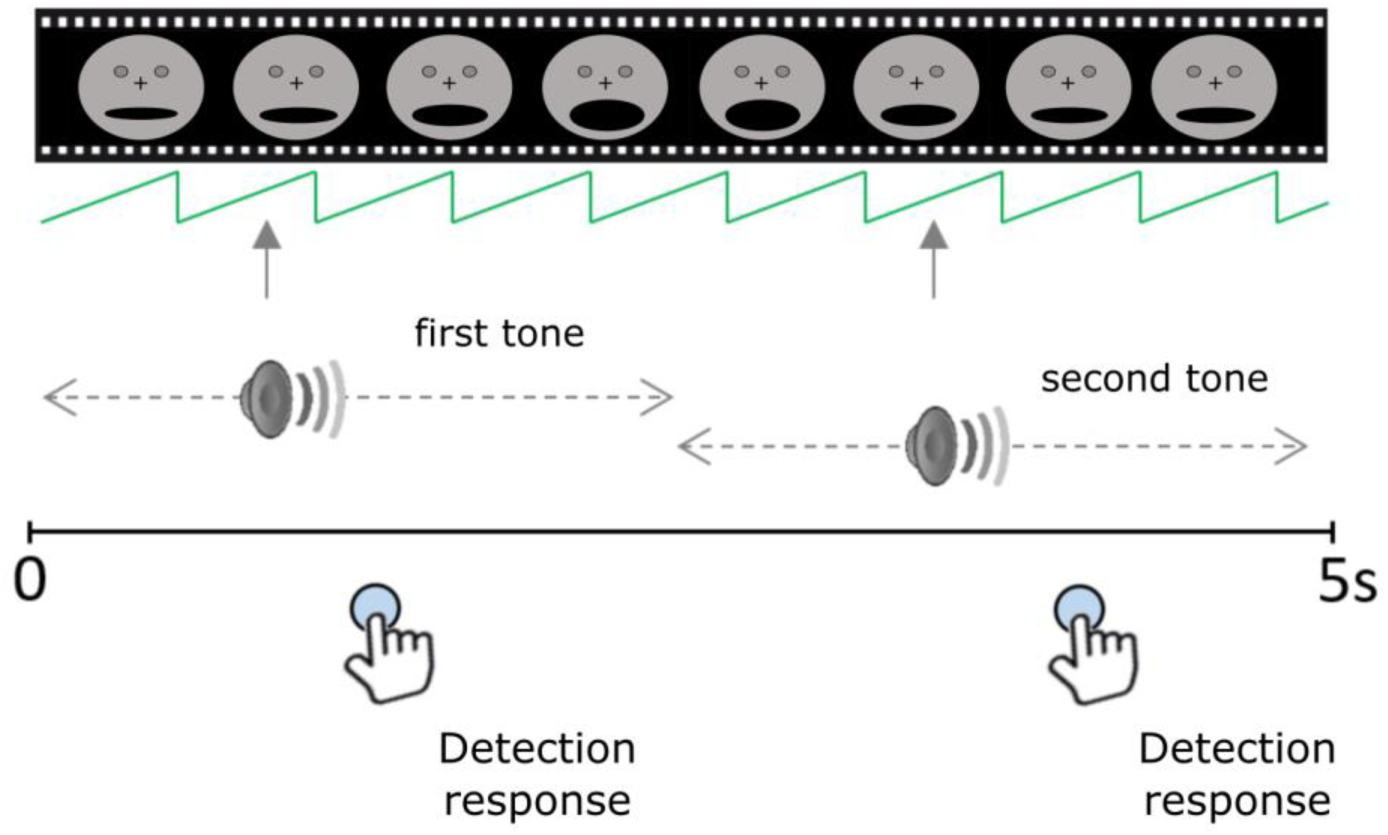
Experimental Paradigm of the Tone Detection Task (TDT). For each trial, continuous white noise and a silent movie were presented together during five seconds. A first pure tone occurred randomly in the first half of the trial while a second tone occurred randomly in the second half of the trial. Participants were instructed to respond as fast and accurately as possible whenever they detected a tone. In the one tone condition, the white noise track contained only one tone that occurred randomly between the two halves of the trial. In the zero tone condition, the sound of the trial contained only white noise (*N.B.* The face of the speaker has been blurred only in the figure for anonymity purpose).

We compared the mean theta phase distributions between first and second tones’ onsets across participants (Figure 2A; see Material and Methods). For each participant, the corresponding theta phases in ongoing lip activity at detected first and second tone onsets were averaged across hit trials. Individual mean theta phases were then averaged across subjects to estimate phase locking of hits to the theta signal conveyed visually in the first and second tone time-windows. Two Rayleigh’s uniformity tests were performed on the first and second grand average theta phase distributions separately. For the first tone window, the Rayleigh’s test did not reject the hypothesis of uniform distribution (n = 24; µ = 1.944 rad or 111.384**°**; r = 0.282; p = 0.148, Bonferroni-corrected). In contrast, the Rayleigh’s test revealed that mean phases were not uniformly distributed in the second tone window (n = 24; µ = −0.999 rad or 302.763**°**; r = 0.44; p < 0.01, Bonferroni-corrected). Further, a permutation test was performed on the resultant vector length (r) difference between the first and second tones to test whether the strength of visual entrainment in the second tone window was significantly stronger than in the first tone window, which indeed was the case (permutations: 10,000; effect size = 0.158; p = 0.015; Figure 2A; see Material and Methods). We performed two additional permutation tests on resultant vector length differences between hits and misses in the first and second tone windows separately to test whether visual entrainment was related to successful auditory processing. No significant difference of vector length was found in the first tone window (permutations: 10,000; effect size = 0.041; p = 0.405), whereas in the second tone window the resultant vector for hits tended strongly to be longer compared to misses (permutations: 10,000; effect size = 0.228; p = 0.052). Following the same approach, we tested the existence of an interaction between visual phasic modulation (hit versus miss) and tone position with a permutation test on the difference of effect size between the first and second tones (i.e. [r_hit_ - r_miss_]_First tone_ vs. [r_hit_ - r_miss_]_Second tone_). Permutations were computed after randomly shuffling the tone position information from original data. Results revealed that the phase modulation between hit and miss trials was significantly greater in the second tone window as compared to first tone window (permutations: 10,000; effect size = 0.187; p = 0.037). Altogether, these results suggest that the visual phase modulation predicted better the detection of second tones and support the hypothesis of visual entrainment shaping auditory perception.

**Figure 2:**
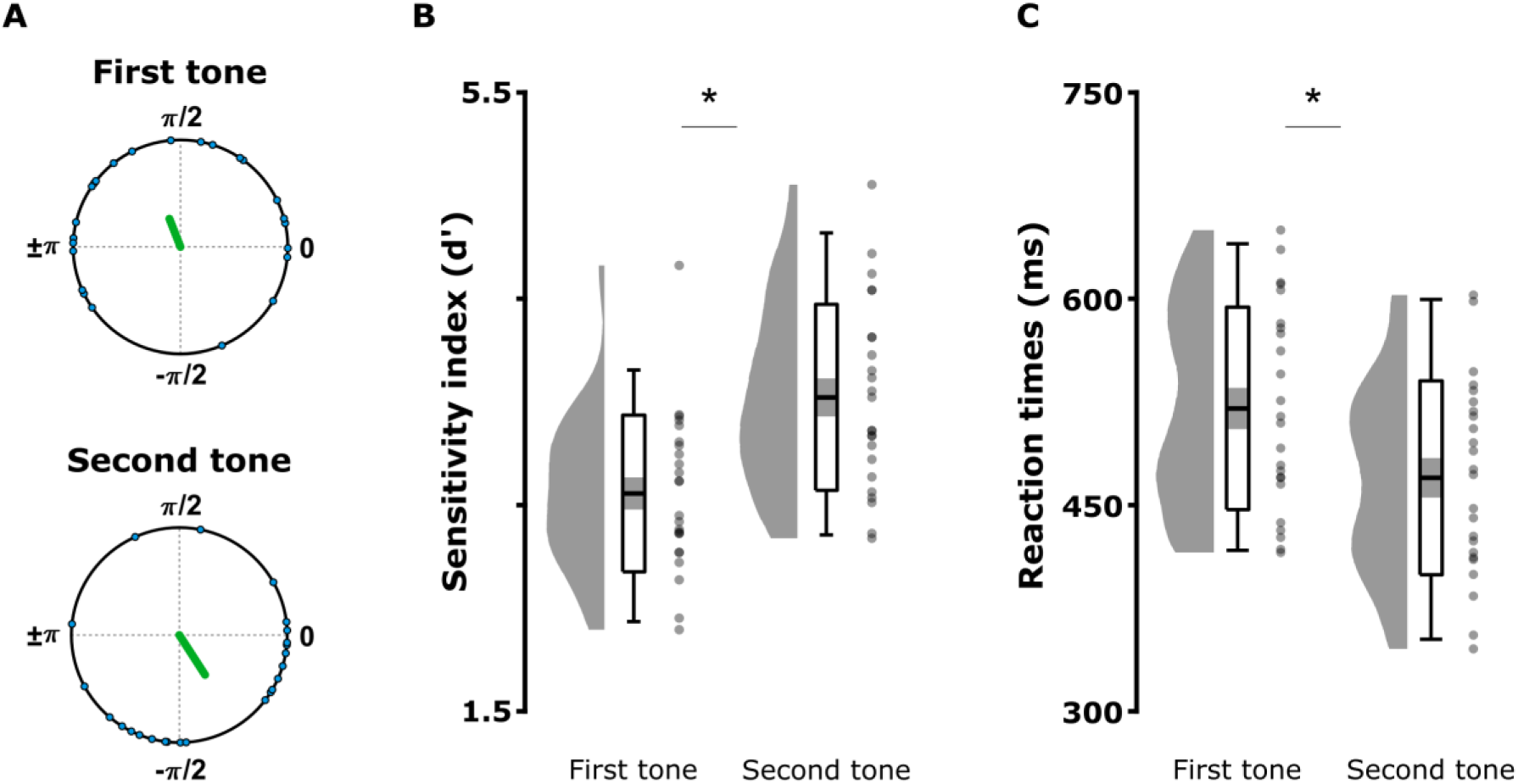
Visual entrainment and tone detection performance in the two tones condition. (A) Resultant r vector length (green line) from grand average phase at the onsets of correctly detected first and second tones across participants. The individual mean theta phases are depicted in polar coordinates (blue circles). (B) Mean sensitivity index (*d’*) and (C) reaction times of first and second tone hits. The graphs depict the density, the grand average (mean ± standard error of the mean; errors bars: 95 % confidence interval), and individual means (grey dots) for first/second tones. Significant contrasts are evidenced with stars (p < 0.05).

Following up, we investigated whether tone detection differed between the first and second tones windows. Such a difference might reflect an auditory bias by visual inputs (Figure 2B). We computed the hit and false alarm (FA) rates to calculate the sensitivity index (*d’* = Z_Hit rate_ - Z_FA rate_). False alarms were calculated by sorting participant’s responses in the absence of a tone, i.e. in the zero tone condition and according to their onsets (occurring either in the first window or in the second window). Because FA rates were very low (Fa_first tone_ = 0.014 ± 0.016; FA_second tone_ = 0.013 ± 0.019), extreme hit and FA rate values were adjusted using a standard *d’* correction by replacing rates of FA rate = 0 with FA rate = *0.5/n*_*noise trials*_ = 0.005 and hit rate = 1 with hit rate = *(n−0.5)/n*_*signal trials*_ = 0.995 (Macmillan & Kaplan, 1985). First, two independent one-sample t-tests established that participants detected the first and second tones in the two tones condition, as the d’ scores were greater than zero (first tone: T(1,23) = 28.02; p < 0.001, two-tailed; second tone: T(1,23) = 28.699; p < 0.001, two-tailed). Second, a paired-samples t-test confirmed that the second tones were better detected than the first ones (T(1, 23) = 5.771 p < 0.001; two-tailed; Fig. 2B). Third, a paired-sample t-test applied on the hit reaction times showed that participants responded faster to second compared to first tones (T(1, 23) = 5.486; p < 0.001; two-tailed; Figure 2C). Therefore, TDT performance difference was driven by hit responses, which were phase modulated as established previously. Importantly, the improvement of the second tone detection could not be attributed to a simple attentional effect due to the presence of the preceding first one, as the single tone condition replicated the two tones condition performances (i.e. by sorting the single tones as first/second tones according to their onsets; see Figure S1B). Finally, a paired-samples t-test performed on the FA rates in the no tone condition confirmed that detection performance modulations did not reflect a change in response bias between the two windows (T(1, 23) = 0.627; p = 0.537; two-tailed). Additionally, the hit rates in both conditions confirmed that the calibration task worked efficiently with mean hit rate = 0.77 ± 0.10 in the two tones condition (first tone = 0.69 ± 0.13; second tone 0.85 ± 0.98) and mean hit rate = 0.74 ± 0.15 in the single tone condition (first tone = 0.67 ± 0.18; second tone = 0.81 ± 0.12). Altogether, these results established that entrainment to theta lips activity increased in time and coincided temporally with increases in auditory detection. In the next step, we aimed at establishing whether a potential audiovisual communication relying on the critical theta organisation of information flows was reflected in the brain.

### Visual cortex leads synchronization to left auditory cortex via theta oscillations during silent lips perception

The above results suggest that visual speech stimuli may recruit the auditory regions via entrainment to render some time-windows more sensible to auditory detection than others. To test this hypothesis on a neural level, we recorded the EEG signal of 23 new participants during the perception of the same 60 silent movies used in the previous tone detection task. Participants were instructed to attend to each movie and rate its emotional content based on the speaker’s face. The movies were presented in a single block and randomised across participants. First, the sources of interest responding to speakers’ lip movements were identified applying a linearly constrained minimum variance beamforming method. Neural entrainment to lip movements was estimated by computing mutual information (MI) on the theta phase between the EEG epochs and corresponding lip signals in the equivalent first tone (0 to 2.5 s; early time-window) and second tone windows (2.5 to 5 s; late time-window). Just as in the behavioural data, we assessed whether entrainment increased over time by contrasting the difference of MI between the early and late time window. Second, the EEG data at the identified visual and auditory sources were reconstructed to perform single-trial phase coupling analysis. The synchrony between visual and auditory sources was reflected by the distribution of theta phase angle differences ϕ_A-V_ = ϕ_audio –_ ϕ_visual_ at each time-point within the early and late time-windows, and the directionality of the coupling was evidenced with the sign of ϕ_A-V_ (i.e. a mean distribution of ϕ_A-V_ = 0 would mean perfect phase alignment, while ϕ_A-V_ < 0 would mean that the visual phase leads the auditory phase; see Material and Methods). To address potential issues of circularity, the directionality of information flow was also assessed using the Phase Slope Index (PSI).

Source localisation analysis revealed that the maximum increases in MI_late_ as compared to MI_early_ were localised in the left visual and auditory cortices, as well as in the right visual cortex to a lesser extent (Figure 3A). This result confirmed the expected recruitment of both visual and auditory sensory areas during the perception of speakers’ lip movements even in the absence of speech sound based on previous studies (Crosse et al., 2019; Calvert et al., 1997). Two separate Rayleigh tests confirmed non-uniform distributions of ϕ_A-V_ in the early (n = 23; µ = −1.99 rad or −114.45°; r = 0.768; p < 0.001, Bonferroni-corrected) and late time-windows (n = 23; µ = −0.92 rad or −52.79°; r = 0.875; p < 0.001, Bonferroni-corrected). An additional Kuiper two-sample test showed that the mean ϕ_A-V_ distributions between the early and late time-windows converged towards two different preferred angles (k = 3.614×10^5^; p < 0.001; Figure 3C). The negative theta phase angle differences ϕ_A-V_ in both the early and late time-windows confirmed that the visual phase led the auditory phase, in line with the idea of visual oscillations responding first to the lips inputs and then directing theta oscillations in the auditory cortex. Additional PSI analyses revealed negative values in both time-windows (respectively PSI_early_ = −0.035 ± 0.055 and PSI_late_ = −0.041 ± 0.035; seed region: auditory source) and confirmed that the left visual source led the left auditory source during silent moving lips perception. Together, these results support our hypothesis that visual cortex led synchronization to left auditory cortex via theta oscillations during silent lips perception. Further, the distance between visual and auditory phases reflects the speed at which information from the visual cortex is conveyed to the auditory cortex. We hypothesized that the phase difference can index the ease of communication between the auditory and visual cortex. A decrease of ϕ_A-V_ over time suggests that visuo-auditory communication gets more efficient, and does not reflect the time lag of a passive transfer of information between the two sensory areas (which in the case would be constant during the entire visual stimulation). To address the latter, we compared the resultant vector length r of the distance between the ϕ_A-V_ phase difference observed in the data and a theoretical ϕ_A-V_ = 0 degree in the early and late time-windows separately. A paired-samples t-test showed that the resultant vector length r of the distance between the observed ϕ_A-V_ and the zero ϕ_A-V_ was significantly greater in the late time-window (T(1, 22) = −2.135; p = 0.044; two-tailed), confirming that coupling between auditory and visual sources became more efficient with time (Figure 3B, C and D).

**Figure 3:**
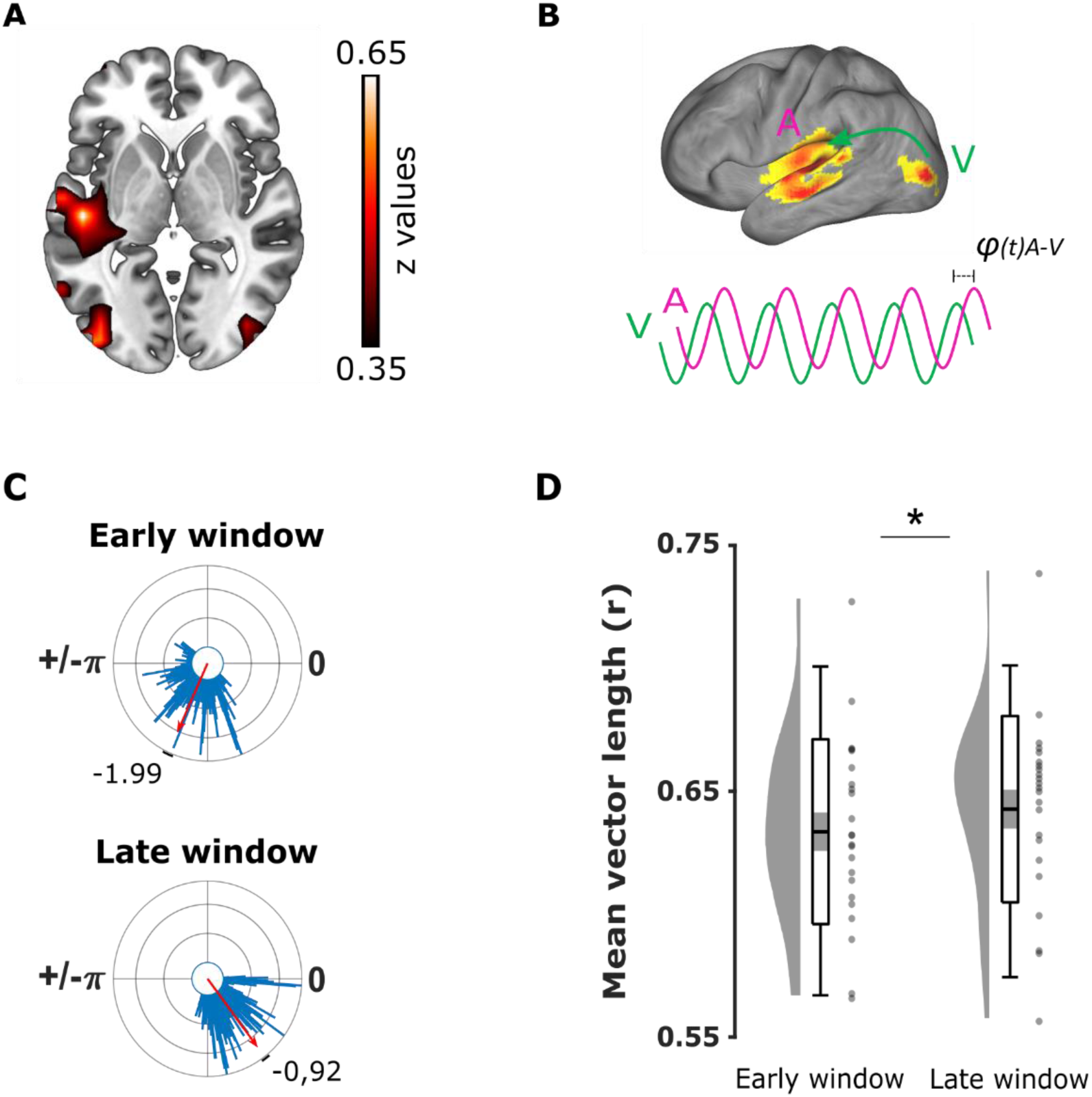
Theta phase coupling analysis between visual and auditory areas during lips perception. (A) Difference of mutual information between the late (2.5 - 5s) and early (0 - 2.5s) time-windows (MI_late_ > MI_early_ contrast; z values). Auditory (MNI coordinates of maximum voxel: [−50 −21 0]; Left Middle Temporal cortex) and visual (MNI coordinates of maximum voxel: [−40 −89 0]; Left Middle Occipital cortex) sources were localized in the left hemisphere. (B) MI_early_ > MI_late_ contrast projected on brain’s surface for illustrative purpose: Synchronisation was estimated through ϕ_A-V_ theta phase offset between theta oscillations at identified auditory (pink line) and visual sources (green line) by mean of phase coupling analysis. (C) Audio-visual phase coupling in the early and late time-windows corresponding to the time-windows containing the first and second tones in the TD Task. The mean ϕ_A-V_ offset between auditory and visual theta phases (red arrows) confirmed that oscillations entrained by lip movements in the visual cortex preceded oscillations in the auditory cortex by 1.99 rad (∼ 56.10 ms) and 0.92 rad (∼ 25.88 ms), respectively in the early and late time-windows. (D) Theta synchronisation between visual and auditory areas improves with entrainment. The resultant vector length r of the distance between the observed ϕ_A-V_ and the theoretical ϕ_A-V_ = 0 was greater in the late than the early window, suggesting a more efficient communication reflected by a decrease of time lag between visual and auditory activities. The graphs depict the density, the grand average (mean ± standard error of the mean; errors bars: 95 % confidence interval), and individual resultant vector length r (grey dots). Significance evidenced with a star (p < 0.05).

## DISCUSSION

In two complementary experiments, we first established that visual entrainment to theta lip phase modulated auditory detection, even if information from silent movies was irrelevant to perform the task. Second, the perception of silent moving lips entrained theta oscillations in the visual cortex followed by the auditory cortex. Together, these results suggest that the brain’s natural reaction to visual speech stimuli might be to align the excitability of the auditory cortex with sharp mouth-openings because that is when one expects to hear corresponding acoustic syllable edges (Hickock & Poeppel, 2007; Giraud & Poeppel, 2012; Peelle & Sommers, 2015 Park et al., 2016; Chandrasekaran et al., 2009). Such a neural process could be a very effective filtering method to increase the sensitivity of the auditory cortex in these relevant time windows for speech comprehension.

Our EEG results suggest that theta oscillations in the left visual cortex encoded the lips’ activity first. Then information travelled to the left auditory cortex via phase coupling to shape its activity. Previous findings reported that the auditory cortex tracks both auditory and visual stimulus dynamics using low-frequency neuronal phase modulation during audiovisual movie perception (Luo, Liu and Poeppel, 2010). Other studies reported that the perception of silent lips also recruited the auditory regions (Bourguignon et al., 2020; Crosse et al., 2015; Calvert et al., 1997). Our findings go beyond by establishing how theta oscillations orchestrate visual and auditory cortices through phase coupling to ensure cross-region communication even in a unimodal condition. Furthermore, it is commonly agreed that entrainment takes several cycles from rhythmic inputs to build up (Doelling et al. 2014; Lakatos et al., 2008; Thut et al., 2011; Zoefel et al. 2018). Behavioural and neural indicators of entrainment reported here consistently increased from the first (i.e. early) to the second (i.e. late) time-window of the trial in both experiments. This supports the idea that we indeed observed neural entrainment to lip movements and sheds light on the functional relevance of visual inputs modulating auditory theta rhythms. As visual onsets naturally lead corresponding auditory onsets by 100-to-300 ms in audiovisual speech (Chandrasekaran et al., 2009; van Wassenhove et al., 2005; Pilling, 2009), visual entrainment to lips may act as a filter by increasing excitability in the auditory cortex to windows containing relevant acoustic features. This hypothesis is corroborated by our phase coupling results where visual theta phase systematically led auditory theta phase during silent movie presentation. This result aligns well with previous findings reported in an intracranial EEG study by Besle and colleagues (2008) investigating evoked potentials between visual and auditory cortices during lip movement perception. The authors showed that lip movements activate the visual motion area (∼140ms after stimulus onset), followed by similar responses in the auditory cortex ∼10 ms later. They hypothesized that direct feedforward processes support audiovisual interactions during moving lip perception via direct projections from the visual cortex to the auditory cortex (Hishida et al., 2003; Cappe and Barone, 2005). Our study goes beyond by showing that both sensory areas communicate within each other via a phase coupling mechanism.

Whether such filtering reflected a direct feedforward modulation from the visual cortex to the auditory cortex or involved top-down modulations remains unclear. Firstly, our results show that the theta phase lag between auditory and visual cortices decreases across time, which speaks against a process that would just passively relay activity from visual to the auditory cortex (as such a passive process should be constant over time). Secondly, additional top-down controls may adjust theta phase synchronization between the sensory cortices. Indeed, higher-level sensorimotor areas also activate during speech perception (Park et al., 2016; 2018; Cognan & Poeppel, 2011; Arnal et al., 2009; Pulvermüller et al., 2005; Wilson et al., 2004). Assaneo & Poeppel (2018) demonstrated recently that activity in the motor and auditory cortices couple at theta rate during syllable perception, correlating with the strength of coupling between speech signal and EEG in the auditory cortex. On the other hand, motor areas play a role in temporal analysis of rhythmic sensory stimulation (Biau & Kotz, 2018; Arnal et al., 2015; Fujioka et al., 2015; Morillon et al., 2019). Entrainment to lip movements may provide the temporal theta structure of speech signal to motor cortex, which in turn adjusts downstream auditory excitability at critical windows containing the corresponding acoustic features in a top-down fashion (in line with Park et al. 2015).

Similarly, Thorne and Debener (2014) associated crossmodal phase resetting with neural temporal predictions occurring when rhythmic visual input precedes auditory input within a stable time-window (∼30 to 100ms). Alternatively, mouth-opening perception may target internal articulatory representations and help to identify the corresponding sounds in the auditory signal. A recent MEG study investigated the visuo-phonological mapping mechanisms mediated by top-down motor areas during silent moving lip presentation (Hauswald et al., 2018). The authors found a stronger coherence between theta activity evoked in the visual cortex by the perception of lip movements and the absent corresponding auditory signal when the moving lips were presented forward as compared to backward. Further, they reported a stronger connectivity between visual cortex and precentral areas in the forward condition, supporting a role of top-down motor controls to map phonological representations with intelligible mouth openings. Here, theta activity found in the auditory cortex may reflect the contribution of inferences generated from such a visuo-phonological mapping. Although speculative, this could partially explain why the increase of entrainment in the auditory cortex was left lateralized, i.e. by recruiting language-related representations classically associated with the left hemisphere. The silent moving lips were taken from speakers speaking in English and presented to native English speakers, which might facilitate such lip-sound mapping. This hypothesis fits with recent debates on whether neural tracking during speech processing reflects online cooperation between pure entrainment to external features and endogenous rhythms providing abstract representations (Meyer, Sun & Martin, 2019; Obleser & Keyser, 2019; Haegens & Zion Golumbic, 2018; Rimmele et al., 2018). However, this would not explain why visual speech information improved the detection of unrelated pure tones here, which will be addressed in future experiments.

The present study focuses on the theta band that reflected the syllabic structure and was dominant in our visual stimuli. Our specific interest towards theta-syllable activity in moving lips was motivated by the fact that the mouth constitutes preponderant direct access to visual speech information mapping sounds (Thompson & Malloy, 2004; Thompson, 1995). However, other parts from the speaker’s face bear quasi-rhythmic speech features at distinct time-scales (Kösem & van Wassenhove, 2017). For instance, eyebrow and head movements temporally align with auditory prosody occurring at 0.5-3Hz delta rate during continuous speech (Ghitza, 2017; Biau et al., 2016; Swerts & Krahmer, 2007; Munhall et al., 2004). Therefore, the proposed filtering method increasing the sensitivity of the auditory cortex may extend to other frequency bands. Future experiments presenting silent videos focusing on speaker’s eyebrows will need to test whether visual entrainment to prosodic features recruits auditory cortex and supports cross-region communication as well; or whether long-range synchronization relies specifically on theta even when the dominant activity in stimuli peaks in other frequency bands.

Finally, additional post-hoc analysis suggested that two subpopulations of participants showed distinct visual theta phases shaping auditory perception in the TDT (Figure S2). One could speculate that participants from the group 2 were fine-tuned to a preferred visual theta phase that represents an optimal time-window, as the effect of tone position on auditory detection was greater than in the group 1. This optimal window allowed information to travel to the auditory cortex either directly or via top-down modulations, and reset auditory activity at “perfect” moments when a tone occurred. Back to our filtering hypothesis, visual theta entrainment would increase auditory excitability coinciding with more time windows containing a tone in this subpopulation regardless of the nature of sounds. During audiovisual speech perception instead, mouth openings may predict critical windows containing envelope peaks relevant for signal processing (i.e., syllable onsets and nuclei). Inversely, mouth closings may loosen up the filtering action when auditory envelope becomes less informative (e.g., silences or noise). Future experiments will determine whether the visual filter tunes auditory activity independently from its relevance, or whether the influence of visual information on auditory activity is restricted to useful time-windows (i.e., only when lip movements predict sound features).

## CONCLUSION

Although the auditory signal alone often provides enough structural information for the early analytic steps of continuous speech, e.g. telephone conversations, a visual filter may be especially helpful to sharpen auditory perception when hearing is impaired or in elders (Grant et al., 1998). Our results provide an important step toward understanding how visual information functionally drives auditory speech perception, and suggest future directions to investigate hearing loss compensation, i.e. to improve lip-reading along with hearing correction.

## MATERIAL AND METHODS

### 1-Tone detection experiment

#### Experimental model and subject details

Twenty-eight healthy English native speakers (mean age = 19 years ± 0.69; 21 females) took part in the first behavioural experiment. Five participants were left-handed. All of them reported normal or corrected-to-normal vision and hearing. All participants were granted experimental participation credit. The data from four participants were excluded because of extreme overall performances and the final analysis were applied on twenty-four data sets. All the participants signed informed consent and ethical approval was granted by the University of Birmingham Research Ethics Committee, complying with the Declaration of Helsinki.

### Method details

#### Apparatus

The task was programmed with Matlab (R2018a; The MathWorks, Natick, MA, USA) and presented with Psychophysics Toolbox (Brainard, 1997; Pelli, 1997; Kleiner et al., 2007). The silent videos were presented on a 21-inch CRT display with a screen refresh rate of 75 Hz (Nvidia Quadro K600 graphics card: 875 MHz graphics clock, 1024 MB dedicated graphics memory; Nvidia, Santa Clara, CA, USA). The auditory stimuli were presented through EEG-compatible insert earphones (ER-3C; Etymotic Research, Elk Grove Village, IL). The accuracy of movie and sound presentation timing was optimised by detecting a small white square displayed on the left of the first frame of each visual stimulus with a photodiode (ThorLabs DET36A, thorlabs.de), and Psychophysics Toolbox (PsychPort Audio and ASIO4ALL extensions for Matlab). Additionally, a parallel audio port was used to record the online audio signal of each trial during presentation. Continuous photodiode and audio data during trials were recorded through a BioSemi Analog Input Box (AIB) adding two separate channel inputs into BioSemi ActiveTwo system: the BioSemi AD-box was connected with the AIB through optical fibres. The input from the photodiode was connected through a BNC connector and the input from the microphone was connected through the 3.5 mm audio. Those two inputs were connected to the AIB though a 37 pin Sub-D connector. Data were digitized using the BioSemi ActiView software, with a sampling rate of 2048 Hz. Offline analysis were performed to calculate the real delay between visual and audio stimuli offset using in-house Matlab codes. Any lag between visual and auditory stimuli onsets was later compensated in the data analyses when computing the corresponding visual theta phase to the tones onsets. The experiment was run from a solid-state hard drive on a Windows 7-based PC (3.40 GHz processor, 16 Gb RAM). Participants used a standard computer keyboard to respond to the task.

### Stimuli

#### Movies

Sixty five-second movies were extracted from natural face-to-face interviews published on YouTube (www.youtube.com) by various universities channels and downloaded via free online application. Satisfying movies containing meaningful content (i.e. one complete sentence, speaker facing toward the camera) were edited using Shotcut (Meltytech, LLC). For each movie, the video and the sound were exported separately (Video: .mp4 format, 1280 x 720 resolution, 25 frame per second, 200 ms linear ramp fade in/out; Audio: .wav format, 44100 Hz sampling rate, mono).

### Lip movements’ detection

Lips contour signal was extracted for each video using in-house Matlab codes. We computed the area information (area contained within the lips contour), the major axis information (horizontal axis within lip contour) and minor axis information (vertical axis within lip contour) as described in Park et al. (2016). In the present study, we used vertical aperture information of the lips contour to establish the theta correspondence between lips and auditory speech (i.e. aperture between the superior and inferior lips) but using area information gave very similar results, as also reported in Park et al. (2016). The lips time-series was resampled at 250 Hz for further analyses with corresponding auditory speech envelope.

### Auditory speech signal

The amplitude envelope of each movie sound was computed using in-house Matlab codes (Park et al., 2018; 2016; Chandrasekaran et al., 2009). First, eight frequency bands equidistant on the cochlear map in the range 100–10,000 Hz were constructed (Smith et al., 2002). Then, sound signals were then band-pass filtered in these bands with a fourth-order Butterworth filter (forward and reverse). Hilbert transform was applied to obtain amplitude envelopes for each band. These signals were then averaged across bands and resulted in a unique wideband amplitude envelope per sound signal. Each final signal was resampled to 250 Hz for further theta correspondence analyses.

### Mutual information between lip movements and corresponding auditory speech signal

To identify the main oscillatory activity conveyed by the lip movements in each visual stimulus, we determined at which theta frequency the auditory and visual speech signals showed significant dependencies. To do so, we examined the audiovisual speech frequency spectrum (1 to 20 Hz) and computed the mutual information (MI) between the minor axis information and speech envelope signals sampled at 250 Hz. MI measures the statistical dependence between two variables with no prior hypothesis, and with a meaningful effect size measured in bits (Ince et al., 2017; Shannon, 1948). We applied the Gaussian Copula Mutual Information (GCMI) approached described in Ince et al. (2017) in which the MI between two signals corresponds to the negative entropy of their joint copula transformed distribution. This method provides a robust, semiparametric lower bound estimator of MI by combining the statistical theory of copulas together with the closed-form solution for the entropy of Gaussian variables, allowing good estimation over circular variables, like phase as well as power. For each movie, the complex spectrum is normalized by its amplitude to obtain a 2D representation of the phase as points lying on the unit circle for both the lip movements and auditory envelope time-series. The real and imaginary parts of the normalized spectrums are rank-normalized separately and the phase dependence for each frequency between the two 2D signals is estimated using the multivariate GCMI estimator giving a lower bound estimate of the MI between the phases of the two signals. Here, we applied the GCMI analyses in two conditions to determine the frequency of interest in each movie: first, we computed MI between corresponding lips and envelope signals as well as non-matching signals (i.e. lips time-series paired with random auditory envelope signals). For the matching signals, the averaged MI spectrum revealed a greater peak in the expected 4 - 8 Hz theta frequencies, reflected by a bump in the band of interest. In contrast, there was no relationship between random auditory and visual signal pairs, which depicts a flat line profile along the whole spectrum (see Supplementary Information Figure S3 A). These results are well in line with previous studies using coherence or MI measures, and confirm the temporal coupling between lips and auditory speech streams at the expected syllable rate in our videos (Park et al., 2016; 2018; Chandrasekaran et al., 2009). Second, for each movie, we performed a peak detection on the MI spectrum to determine which specific frequency carried most theta information to maximize entrainment in the tone detection and silent movie perception tasks (4Hz frequency peak: 16 videos; 5Hz frequency peak: 15 videos; 6Hz frequency peak: 9 videos; 7Hz frequency peak: 13 videos; 8Hz frequency peak: 7 videos. See Supplementary Information Figure S3 B).

### Audio tones and white noise

Pure auditory tones and white noise stimuli were generated using in-house Matlab codes. The target tone consisted in a sinusoidal signal of 100 ms at one kHz (sampling rate: 44100 Hz). The same noise consisted in a Gaussian white noise lasting two seconds for the calibration task and five seconds in the tone detection task (the white noise has been generated only once and loaded during each procedure to ensure that all the participants were tested with the same noise; sampling rate: 44100 Hz). Both the tone and the white noise signals were normalized between - 1 to 1 (arbitrary units) and the loudness of the resultant stimuli (white noise with embedded tone) was estimated at ∼ 91 dB SPL sound pressure level (SPL) using in-house Matlab codes (Moore et al., 2016). During the entire procedure, the auditory stimuli were displayed at constant ∼72 dB SPL and across participants.

### Tones onsets

For each trial, the target tones were embodied in the white noise at predetermined pseudo-random onsets counterbalanced across conditions (zero, one or two tones per trial, 100 trials per condition). In the calibration task serving to determine the individual threshold of target tones detection (see below for the general procedure), there could be only zero or one tone maximum per trial. For the one tone condition, the onset of the target tone always randomly occurred between 300 and 1400 ms after the trial onset to allow participants to detect it properly and have time to respond before the end of the trial. In the zero tone condition, the auditory track consisted in two seconds of white noise only. In the tone detection task, there could be zero, one or two tones per trial. In the one tone condition, the onset of the target always occurred randomly between 300 and 4500 ms after the trial onset. In the two tones condition, the first tone randomly occurred in a time-window centred on the first half of the trial length, between 300 and 3000 ms (mean first tone onsets = 1.68 ± 0.78 s). The second tone occurred in a time-window centred on the second half of the trial length, between 1000 ms after the first tone onset and 4500 ms after the trial onset (mean second tone onsets = 3.45 ± 0.62 s). This design provided participants with enough time to detect and respond to both tones, and kept the two tones temporally unrelated from each other. In the zero tone condition, the auditory track consisted in five-second of white noise only. The signal-to-noise ratio between target tones and white noise was determined for each participant individually with the calibration task performances and adjusted consequently in the following tone detection task (see below).

### Procedure of the calibration task and tone detection task (TDT)

The experiment began after the completion of a safety-screening questionnaire and the provision of informed consent. Participants sat in a well-lit testing room at approximatively 60 cm from the centre of the screen and wore the insert earphones for sound presentation. Participants performed first a short pure tone detection task with no visual stimuli (i.e. calibration task). This task served to determine the individual threshold at which each participant detected ∼ 70 - 80 % of the target tones in auditory modality only, and the signal-to-noise ratio (SNR) to be implemented between the amplitude of the target tones and the white noise in the following tone detection task (TDT). The calibration task was composed of a four-trial practice to identify the target tone itself, followed by five blocks containing 20 trials each. Each trial began with a black fixation cross (500 - 1000 ms duration, jittered) followed by the presentation of a red cross over a grey background during two seconds to indicate the period of possible target tones occurrence. A continuous white noise was displayed during the red cross presentation. In 50 % of the trials, a unique audio tone was embedded in the white noise at unpredictable onset, and participants had to press “1” key as fast and accurately as possible only when they perceived a target tone. The pseudo-random sequence of the procedure ensured that there were never more than two consecutive trials of the same condition. The participants received no feedback and the procedure continued to the next trial after the end of the two-second white noise. The signal-to-noise ratio was adjusted following an adapted “two-down one-up*”* staircase procedure (see Leek, 2001): For the first five trials, the SNR was fixed (mean white noise power of 0.981) and served as a starting point across participants. After each trial, the keypress response of the participant was stored to adjust the SNR for the next trial as following: for two successive hits, the SNR was decreased by 2 % of the starting signal energy in the next trial. For two successive correct rejections (i.e. no response when no tone occurred) or one correct rejection following a hit, the SNR was kept identical for the next trial. After a miss or a false alarm, the SNR was always increased by 2 % of the starting signal energy. At the end of the calibration task, the individual SNR was averaged over the last 30 trials and stored for the following real tone detection task (mean calibration accuracy rate: 0.75 ± 0.05). The participants took a short break and were recalled the instructions before starting the proper tone detection task. The calibration task lasted approximatively seven minutes.

The main structure of the TDT was the same as in the precedent calibration task. The TDT was composed of a short four-trial practiced followed by 300 trials divided in 6 blocks of 50 trials each and separated by breaks (the sixty silent movies were repeated five times each to generate the total 300 trials). Each trial began with a red fixation cross presentation (500-1250 ms duration, jittered). Then, a random five-second silent movie was presented with a black fixation cross in the centre of the screen to give the participants a point to gaze at and reduce saccades. The continuous white noise was displayed together with the silent movie according to the three random conditions: no tone (100 trials), one single tone (100 trials) or two tones (100 trials) hidden in the white noise. Participants were instructed to press “1” key as fast and accurately as possible only when they perceived a target tone. The participants received no feedback on their responses and the procedure continued with the next trial after the end of the silent movie. The SNR between the tones and the white noise was determined in the previous calibration task as explained above. The TDT lasted approximatively 50 minutes.

### TDT conditions

The condition of interest containing the two tones (i.e. first and second tone) served to assess our main hypothesis that entrainment increases in time with the perception of visual information conveyed by the speakers’ lip movements. According to this, the second tones should be better detected and associated to a greater theta entrainment as compared to the first tones to reflect the direction of the auditory system by the entrained visual system to lip movements. The zero and single tone conditions were additional control conditions: the zero tone condition served to determine the false alarm rates (i.e. participants’ keypresses in the absence of tone) and controlled whether participants tended to press more together with the tone onset delays (i.e. time-dependent response bias). The single tone condition served to counterbalance the number of trials containing two tones and control for the predictability of the second tone. The replication of the performances observed in the two tones condition by sorting the single tones according to their onsets equivalent to either first or second tone onsets would confirm that the detection of the second tone is not due to its predictability from a preceding tone but its position in time only. The pseudo-random sequence of the procedure ensured that there were never more than three consecutive trials of the same condition.

### Quantification and statistical analysis

The Tone Detection task was within-subject design.

### Tone detection performances

The hits (i.e. correctly detected tones) and false alarms (i.e. keypress responses during the zero tone condition allocated to the first or second tone windows depending on their onsets) rates were computed to calculate the individual mean sensitivity index (i.e. d’) in the two conditions for each participant (i.e. single tone and two tones conditions). The reaction times of the hits were computed to calculate the individual mean reaction times in the two conditions for each participant (i.e. single tone and two tones conditions). Additionally, we calculated the mean correct response rates and reaction times of the two conditions concatenated together of each individual to exclude blindly potential outliers without favouring the results towards our hypothesis and performing as following: below chance level (correct response rate < 0.5) or perfectly (correct response rate = 1), or with mean reaction times outside the grand averaged reaction times ± two standard deviations range. Accordingly, four participants were excluded from analyses (two participants performed below chance level, one participant performed perfectly and one participant’s reaction times were slower than the grand average mean + two standard deviations). A paired-samples t-test was conducted on the averaged d’ scores and hit reaction times between the first and second tones in the two tones condition and single tone condition separately. Additionally, a paired-samples t-test was performed on false alarm rates from the first and second windows in the zero tone condition to control for any response bias with time.

### Visual entrainment to theta activity conveyed by lip movements

To bridge visual entrainment to auditory processing together, we related the tone target onsets to the theta activity conveyed by the lip movements during silent movies perception: First, for each movie the theta phase of the lip movements’ time-series was computed by applying a Hilbert transform with a bandpass filter centred on the frequency bin of MI peak ± 2 Hz, accordingly to the mean theta frequency determined in MI stimuli analyses. Second, we computed the instantaneous theta phase of the lips signal corresponding to the onset of the tones occurring during each trial. All further circular statistics on angular scale were performed using the CircStat toolbox on Matlab (Berens, 2009). The circular uniformity in the first and second tones windows within and across participants were estimated separately by applying Rayleigh tests to calculate the mean direction and resultant vector length from hits/miss trials. To assess statistically the strength of visual entrainment between the first and second tone windows in the two tones condition (hits only), we performed a permutation test on the resultant vector length difference (z-value) second tone minus first tone reflecting the effect size. For each participant, we generated 10000 iterations as following: first, the hit trial labels were shuffled between the first and second tones in the two tones condition. Second, two balanced subsamples of shuffled trials were selected, with a number matching the smallest number of trials between the first and second tone hits. Third, the mean phase of the first and second tone shuffled trials were computed for each iteration and per participant. Fourth, a Rayleigh’s test of uniformity was applied on the mean phases to determine a resultant vector length at the first and second tones per iteration (i.e. z-value). For each iteration, we computed the difference of z-value_second tone_ - z-value_first tone_ to quantify its effect size, and the resultant 10000 z-value differences were sorted in descending order. To estimate the final p-value and test the null hypothesis, the difference of z-value between the original first and second tone data was ranked in the sorted permuted z-value differences and divided by the total number of permutations+1. If the p-value was smaller than α = 0.05, we rejected the null hypothesis H_0_ = there is no difference of resultant vector length between the first and second tones (i.e. the visual entrainment is significantly greater in the second tone window). The exact same approach was applied on the single tone condition, as well as on the hit versus miss entrainment comparisons.

### Subpopulations of group 1 and group 2

Participants were sorted in two subpopulations according to their preferred theta phase in the second tone window, were visual entrainment supposedly took place after enough lip movements inputs in the condition of interest (i.e. two tones condition). The Rayleigh tests revealed non-uniform distributions of preferred phase at the second tones for group 1 (n = 11; µ = 348.83°; p < 0.001) and group 2 (n = 10; µ = 249.63**°**; p < 0.001). A Kuiper two-sample test confirmed that the mean preferred phases were different between group 1 and 2 (k = 121; p < 0.01). Results are presented in the supplementary information.

### 2-Silent movie perception-EEG experiment

#### Experimental model and subject details

Twenty-five healthy English native speakers (mean age = 21.52 years ± 3.86; 17 females) took part in the first behavioural experiment. All of them reported normal or corrected-to-normal vision and hearing, and were right-handed. Twenty participants were granted credits and five participants received financial compensation for their participation (£20). The data from two participants were excluded from the final analyses due to too noisy EEG data. All the participants signed informed consent and ethical approval was granted by the University of Birmingham Research Ethics Committee, complying with the Declaration of Helsinki.

### Method details

#### Apparatus

The task was programmed with Matlab (R2018a; The MathWorks, Natick, MA, USA) and presented with Psychophysics Toolbox (Brainard, 1997; Pelli, 1997; Kleiner et al., 2007). The silent videos were presented in the same manner as described in the previous TDT section.

### Stimuli

#### Movies

The movies presented during the silent lips perception task were the exact same 60 movies used in the previous tone detection task. The order of movies was randomized across participants.

#### Procedure

Participants sat in a well-lit testing room at approximatively 60 cm from the centre of the screen to complete a safety-screening questionnaire and the provision of informed consent first. After the correct preparation of the EEG cap, the participants were instructed to attend to all the movies quietly and to avoid movements during the presentation. Each trial was preceded by a central fixation cross (500 - 1250 ms duration, jittered) followed by the presentation of a random five-second movie. A central fixation cross was displayed during the movie presentation to give participants a point to gaze at and reduce excessive saccades. Participants were instructed to attend to each movie carefully and rate its emotional content based on speaker’s facial gestures by using the number keys on the keyboard after the presentation (i.e. 1 for neutral through 5 for very emotional; results not reported). The total presentation of the sixty movies lasted approximatively 10 minutes.

Online EEG recordings: Continuous EEG signal was recorded using a 128 channel BioSemi ActiveTwo system (BioSemi, Amsterdam, Netherlands). Vertical and horizontal eye movements were recorded from additional electrodes placed approximatively one cm to the left of the left eye, one cm to the right of the right eye, and one cm below the left eye. Online EEG signals were digitalized using BioSemi ActiView software at a sampling rate of 2048 Hz. For each participant, the position of the electrodes on the scalp were tracked using a Polhemus FASTRAK device (Colchester) and recorded with Brainstorm (Tadel et al., 2011) implemented in MATLAB.

Offline EEG preprocessing: EEG data were preprocessed offline using Fieldtrip (Oostenveld et al., 2011) and SPM 8 toolboxes (Wellcome Trust Centre for Neuroimaging). Continuous EEG signals were bandpass filtered between one and 100 Hz and bandstop filtered (48–52 Hz and 98–102 Hz) to remove line noise at 50 and 100 Hz. Data were epoched from 2000 ms before stimulus onset to 7000 ms after stimulus onset, and downsampled to 512 Hz. Bad trials and channels with artefacts were excluded by visual inspection and numerical criteria (e.g., variance as well as kurtosis) before applying an independent component analysis (ICA) to remove components related to ocular artefacts. Bad channels were then interpolated using the method of triangulation of nearest. After re-referencing the data to average reference, trials with artefacts were manually rejected by a last visual inspection. On average, 4.48 ± 2.48 trials were removed and 4.04 ± 1.82 channels were interpolated per participants.

#### Head models

For the 22 participants without individual MRI scans, the MNI-MRI and the volume conduction templates provided by Fieldtrip were used to construct the head models. Electrode positions of each participant were aligned to the template head model. Source models were prepared with the template volume conduction model and the aligned individuals’ electrode positions following standard procedures. One participant provided his own MRI scans and his head model was built using his structural scans (Michelmann et al., 2016): the MRI scans were segmented into four layers (i.e. brain, CSF, skull and scalp) using the Statistical Parametric Mapping 8 (SPM8; http://www.fil.ion.ucl.ac.uk/spm) and Huang toolboxes (Huang et al., 2013). The volume conduction model was constructed using the dipoli method implemented in Fieldtrip. Participant’s electrode positions were aligned to his individual head model. Finally, his MRI was warped into the same MNI template MRI of Fieldtrip and the inverse of the warp was applied to a template dipole grid to have each grid point position in the same normalized MNI space as the other participants for further group analyses.

#### Source localization during silent movie perception

Source analyses on EEG data recorded during silent movies presentation were run using individual electrode positions, grid positions and template volume conduction model. For the participant who had his MRI scans, source analyses were calculated using normalized grid positions instead. Source activity was reconstructed using a linearly constrained minimum variance beamforming method implemented in Fieldtrip (LCMV; see Van Veen et al., 1997). The neural entrainment to lip movements at source level was determined by computing mutual information between EEG epochs and the lip movements during silent movie presentation (i.e. lips time-series from the silent movie presented during the trial). To test our hypothesis that entrainment builds up in time with perceived theta lips activity, we contrasted the difference of MI between the equivalent time-window to the second tone window (MI_late_), and the equivalent time-window to the first tone window (MI_early_) in the previous TDT. Accordingly, we expected first to observe an increase of theta activity in the visual cortex reflecting entrainment to lip movements. Second, we expected an equivalent pattern in the auditory correlates reflecting a tuning from visual activity. For each single trial, MI was first computed separately in the time-windows equivalent to the first (0 to 2.5 seconds after trial onset; MI_early_) and second tone time-windows of the TDT (2.5 to 5 seconds after trial onset; MI_late_) at the 2020 virtual electrodes by using the same approach described in the stimuli analysis section (i.e. where we established which frequency carried most correspondence between lips and envelope signals for each video; using a wavelet transform to compute the phase). Second, for each single trial, the MI spectrum was realigned respect to the frequency bin (± 2 Hz) corresponding to the peak of MI between lips and envelope signal established in the movie analyses. This step was done to be able to average all the trials together taking into account the main theta activity carried in each individual movie. For instance, if the peak of MI between lip movements and auditory envelope was found at 4 Hz in the video number 1, the realigned MI spectrum between EEG and lips signals from the trials presenting video number 1 was now 4 ± 2 Hz (2 to 6 Hz; 1 Hz bin) to insure that the central bin of each single trial corresponds to the objectively determined frequency peak of theta activity. Third, the realigned MIs of single trials were averaged across trials within each participant for further group analyses. For each participant, we calculated the normalized difference of MI at the frequency bin of interest in the late minus early time-window at all the 2020 virtual electrodes (MI normalization: (MI_late_ - MI_early_)/ MI_early_; third bin in the realigned MI spectrum). Finally, the normalized difference of MI between second and first tone time-windows was grand averaged across participants and the grand average was interpolated to the MNI MRI template. The coordinates for auditory and visual sources of interest were determined by finding the maximum of MI_late_ - MI_early_ differences in regions corresponding to the auditory and visual areas, and defined using the automated anatomical labelling atlas (AAL, Tzourio-Mazoyer et al., 2002).

#### Source reconstruction

We performed time-series reconstruction analysis to investigate the synchronization at theta activity between the two sources of interest during silent movie presentation. The time series data were reconstructed and extracted at the visual and auditory coordinates determined by source localization analysis. LCMV beamformer reconstruction can cause random direction of source dipoles and eventually affect phase analysis results. To get around this issue, the event-related potentials (ERP) time-locked to movie onsets at visual and auditory sources were plotted to identify the visual component, i.e. N1-P2-N2 waveform (Wang et al., 2018). After visual inspection, the sign of the reconstructed data were flipped in direction by multiplying the time-series by −1 if any visual or auditory source ERPs showed the opposite of the expected direction of a visual component (i.e. negative-positive-negative polarity). This “flipping” correction was applied consistently across all trials before sorting data between early and late time-windows, thus it did not bias results towards our hypothesis. The same phase coupling analyses were computed with unflipped source data as a control. Phase angle differences between visual and auditory theta activities in the early and late windows were also non-uniformly distributed according to Rayleigh tests with significantly different mean angles according to a Kuiper’s test, confirming that the flipping procedure only better reflected phase coupling modulation with entrainment.

#### Theta phase coupling between auditory and visual sources

First, auditory signal was projected orthogonally onto the visual signal applying a Gram-Schmidt process (GSP; Hipp et al., 2012) for single trials before computing phase information. This was done to reduce the noise correlation patterns reflecting activity from a common source (i.e. volume conduction) estimate captured at different electrodes (in that case, the phase alignment reflects the same source activity and not the phase coupling between two distinct source activities). The GSP increases the signal-to-noise ratio by leaving intact the proper activities conveyed at the two distinct electrodes while reducing noise correlation weight (see Hipp et al., 2012). Second, for each trial the instantaneous theta phase of the auditory and visual orthogonalized time-series were computed by applying a Hilbert transform with a bandpass filter centred on the frequency bin of MI peak ± 2 Hz, accordingly to the mean theta frequency of the video presented during the trial. Third, the difference of unwrapped instantaneous phase between auditory and visual sources was computed for each single trial at each time-point in two windows corresponding to the time-windows containing the first (0.5 to 2 seconds after trial onset; early window) and second tones in the TDT task (3 to 4.5 seconds after trial onset; late window). The first and last 500 ms at the edges of the epoch were not included into phase coupling analyses to avoid the trial onset and offset responses. To further control for any potential issues of circularity, the phase-slope index (PSI) was calculated in the early and late time-windows between the left auditory and the left visual sources to assess the directionality of information flow between our two sources of interest (Nolte et al., 2008). The PSI was computed separately in the early and late windows with a bandpass filter centred on the frequency bin of MI peak ± 2 Hz of each single trial, using the left auditory source as the seed region. A positive PSI indicates that the seed region leads the visual source, while a negative PSI indicates that the visual source leads the auditory source during the visual perception of silent moving lips.

#### Quantification and statistical analysis

The silent lips perception task were within-subject design.

#### Audio-visual theta synchrony

The phase coupling between auditory and visual sources was estimated through their theta phase angle difference in time-windows equivalent to the first and second tone windows from the TDT (i.e. MI_early_ and MI_late_ time-windows). To assess that the ϕ_A-V_ theta phase coupling between visual and auditory cortices was modulated in time (i.e. between early and late time-windows), we computed and compared the resultant vector length r of the distance between the observed ϕ_A-V_ in the data and a theoretical angle ϕ_A-V_ = 0 degree in the early and late time-windows separately. We hypothesized that the ϕ_A-V_ reflects the speed at which information from the visual cortex is conveyed to the auditory cortex. If the ϕ_A-V_ is constant over time, this means that it reflects the time lag of a passive transfer of information between the two sensory areas. In contrast, a decrease of theta phase distance over time (i.e. the ϕ_A-V_ in the late time-window gets closer to zero degree than the ϕ_A-V_ in the early time-window) suggests that visuo-auditory communication gets more efficient. For each trial, we calculated the resultant vector length of the distance between the real auditory-visual phase offset and an arbitrary fixed phase offset at 0° at each time point in the two time-windows. The resultant vector length was collapsed across time in the first and second windows separately, resulting in two values per trial. Single-trial values in the first and second windows were then averaged across trials for each participant, and the difference of phase entrainment values was assessed with a paired samples t-test.

## ADDITIONAL INFORMATION

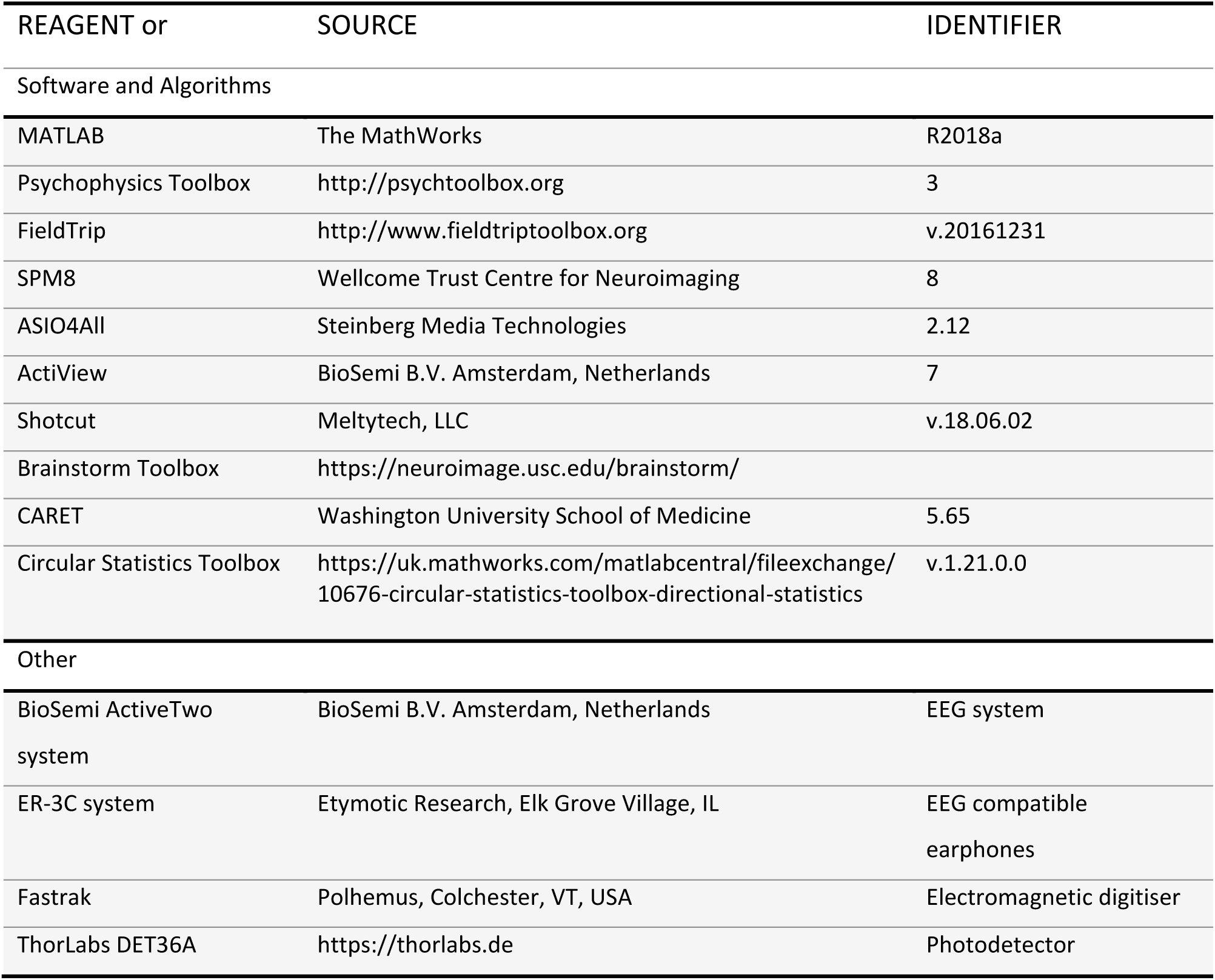

| REAGENT or | SOURCE | IDENTIFIER |
| --- | --- | --- |
| Software and Algorithms |  |  |
| MATLAB | The MathWorks | R2018a |
| Psychophysics Toolbox | http://psychtoolbox.org | 3 |
| FieldTrip | http://www.fieldtriptoolbox.org | v.20161231 |
| SPM8 | Wellcome Trust Centre for Neuroimaging | 8 |
| ASIO4All | Steinberg Media Technologies | 2.12 |
| ActiView | BioSemi B.V. Amsterdam, Netherlands | 7 |
| Shotcut | Meltytech, LLC | v.18.06.02 |
| Brainstorm Toolbox | https://neuroimage.usc.edu/brainstorm/ |  |
| CARET | Washington University School of Medicine | 5.65 |
| Circular Statistics Toolbox | https://uk.mathworks.com/matlabcentral/fileexchange/10676-circular-statistics-toolbox-directional-statistics | v.1.21.0.0 |
| Other |  |  |
| BioSemi ActiveTwo system | BioSemi B.V. Amsterdam, Netherlands | EEG system |
| ER-3C system | Etymotic Research, Elk Grove Village, IL | EEG compatible earphones |
| Fastrak | Polhemus, Colchester, VT, USA | Electromagnetic digitiser |
| ThorLabs DET36A | https://thorlabs.de | Photodetector |

## CONTACT FOR REAGENT AND RESOURCE SHARING

Data for individual participants are available, as consent for sharing data at the level of the individual participant was received. Further information or requests should be directed to the corresponding author.

## AKNOWLEDGMENTS

This work was supported by a Sir Henry Wellcome Postdoctoral Fellowship awarded to E.B (Grant reference number: 210924/Z/18/Z), as well as grants from the ERC (Consolidator Grant 647954) and ESRC (ES/R010072/1) awarded to S.H, who is further supported by the Wolfson Foundation and Royal Society. The authors would like to thank people from the Memory and Attention Lab and David Poeppel for their valuable comments and inputs during the preparation of this manuscript.

## AUTHOR CONTRIBUTION

E.B, H.P and S.H designed the experiments and paradigms. E.B and D.W collected and analysed the data. E.B, D.W, H.P, O.J and S.H wrote the paper. All the authors discussed the results and commented on the manuscript.

## SUPPLEMENTARY INFORMATION

Supplemental Information includes three figures, one silent movie (Movie1) and one white noise containing two tones (Sound1) used in the TDT and EEG tasks.

## SUPPLEMENTARY INFORMATION

### Tone detection performance and visual entrainment in the single tone condition

**Figure S1:**
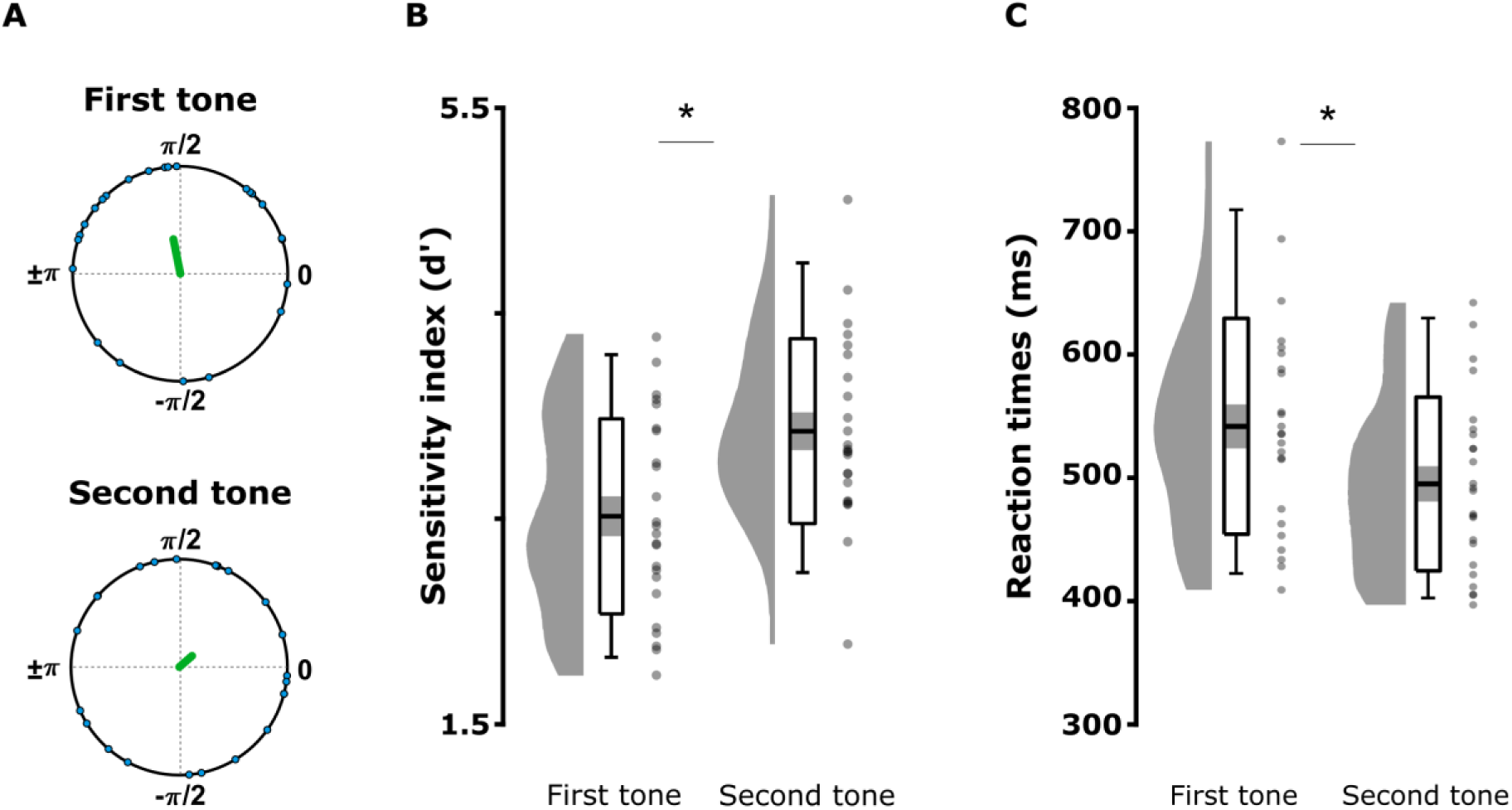
Visual entrainment and tone detection performance in the single tone condition. Although in this condition there was only one tone per trial, we sorted each single tone as a first or second tone according to its onset. (A) Resultant r vector length (green line) from grand average phase at the onsets of first tones hits and second tones hits. The individual mean theta phases are depicted in polar coordinates (blue circles). (B) Mean sensitivity index (d’) and (C) reaction times. The graphs depict the density, the grand average (mean ± standard error of the mean; errors bars: 95 % confidence interval) and individual means (grey dots) for tones sorted as first or second ones. Significant contrasts are evidenced with stars.

To control that the better detection of the second tone did not simply reflect an attentional effect driven by the occurrence of the first tone in the two tones condition (Figure 2B), we compared performance with the single tone condition by sorting each unique tone of the trial as a first or second tone according to its onset (i.e. respectively between 0 and 2.5 s or 2.5 and 5s after trial onset; Figure S1 B&C). Two independent one-sample t-tests established that participants detected the first and second tones in the single tone condition, with d’ scores greater than zero (first tone: T(1,23) = 21.99; p < 0.001, two-tailed; second tone: T(1,23) = 27.69; p < 0.001, two-tailed). Two paired-samples t-test performed on d’ scores and reaction times confirmed that the second tones were better detected (figure S1 B; T(1, 23) = 5.385; p < 0.001; two-tailed), and faster as compared to the first tones (figure S1 C; T(1, 23) = 4.778; p < 0.001; two-tailed). Further, we compared directly the tone detection performances between the single tone and two tones conditions by mean of 2-by-2 repeated-measures ANOVAs (factors condition and tone position). A main effect of position showed that the second tones were better detected than the first tones in both conditions (F(1, 23) = 37.123; p < 0.001). No main effect of condition (F(1, 23) = 2.302; p = 0.143) or interaction between condition and tone position on d’ were found (F(1, 23) = 0.658; p = 0.426). A repeated-measures ANOVA on reaction times showed a significant effect of tone position with faster responses to second tones than first tones (F(1, 23) = 33.797; p < 0.001). A significant effect of condition showed overall faster reaction times in the two tones condition as compared to the single tone condition (F(1, 23) = 14.047; p = 0.001), but no interaction between tone position and condition (F(1, 23) = 0.173; p = 0.682).

Finally, although initially designed to control for attentional bias on detection performance (and counterbalance the number of trials followed by a second tone to control for its predictability), we tested whether the phase modulation reported at the second tone in the two tones condition would be true also at the second tones in the single tone condition as well (Figure S1 A). Similarly to Figure 2A, we compared the resultant r vector length (green line) from grand average phase at the onsets of tones sorted as first tones hits (n = 24; µ = 1.769 rad or 101.336°; r_first_ = 0.329; p = 0.074) and second tones hits (n = 24; µ = 0.719 rad or 41.165°; r_second_ = 0.157; p = 0.558) in the single tone condition. A permutation test on the resultant vector length difference between the tones sorted as first and second tones did not reveal significant difference (figure S1 A; permutations: 10000; effect size = −0.171; p = 0.835). Finally, the permutation test applied to assess the difference of effect size between the first and second tones did not reveal any significant interaction between visual phase modulation and tone detection (permutations: 10000; effect size = −0.172; p = 0.866). The absence of phase modulation here may be explained by the fact that single tones were sorted as first or second tones according to their onset, leaving only half as many trials as in the two tones condition of interest.

### Distinct preferred phases showed performance differences between two subpopulations of listeners

The behavioural data of the TDT task suggests that two separate subpopulations entrained to different preferred theta phases in the second tone time-window (see Figure 2A lower panel and Figure S2B below). In a post-hoc analysis, we assessed whether these apparently distinct populations also showed differences in tone detection performances. Arguably, any difference should be most pronounced only when visual entrainment eventually took place (second tone window) but not early in the trial. Participants were sorted in two groups based on their mean theta phase in the second tone window (i.e. n_group1_ = 11 and n_group2_ = 10; see Material and Methods) and we compared detection performance (d’) in the two tones condition by means of a repeated-measures ANOVA (with factors tone position and group).

**Figure S2:**
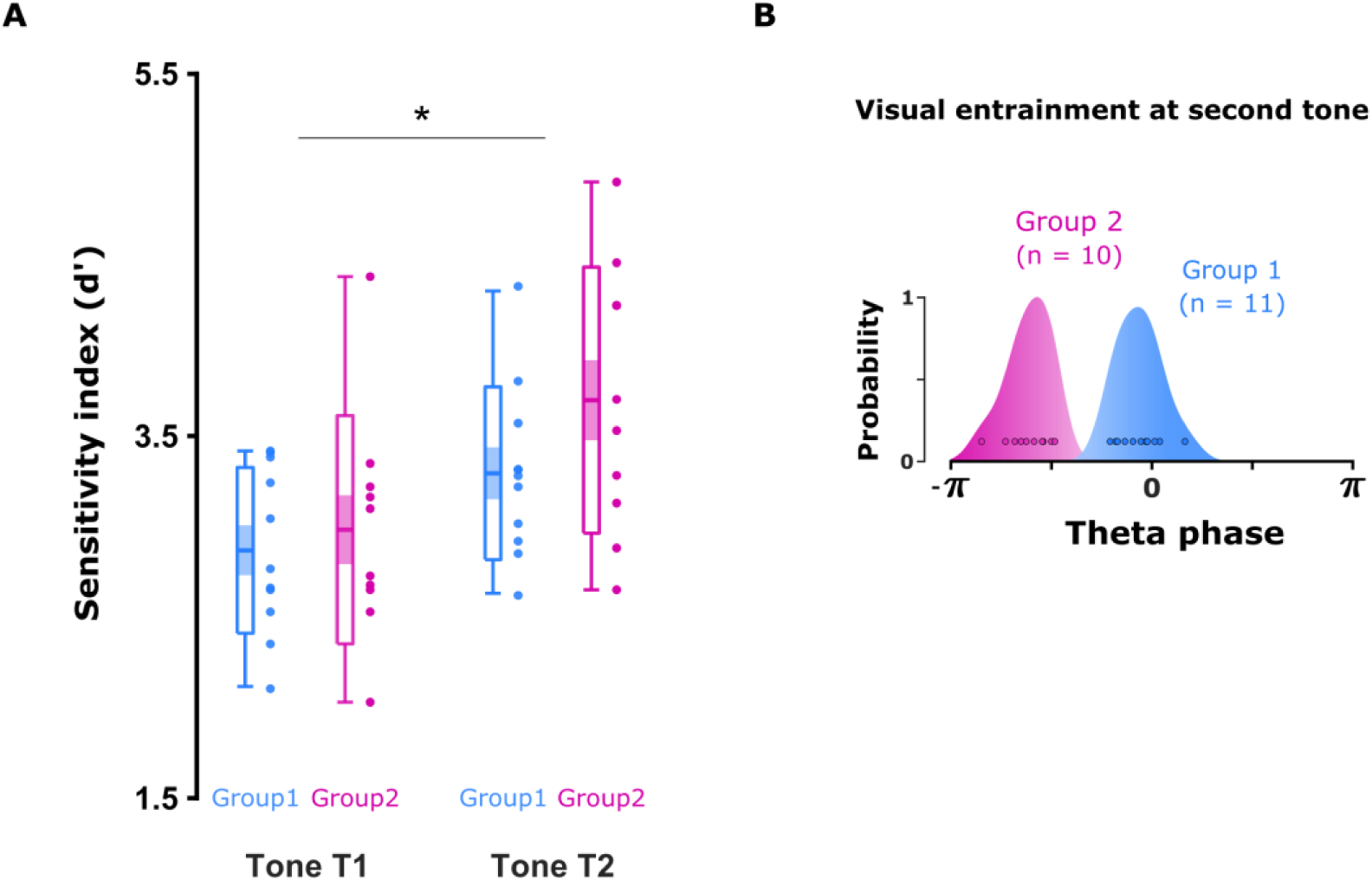
Preferred theta phase affects tone detection task performance. (A) Tone detection sensitivity (d’) of first and second tones between the group 1 and 2. The graphs depict the density, the grand average (mean ± standard error of the mean; errors bars: 95 % confidence interval), and individual means for the first/second tones. Post-hoc analyses revealed that the effect of tone position on the detection performances (first tone versus second tone) was greater in the group 2 than group 1, suggesting that visual entrainment affected auditory processing differently between the two groups. Significant contrasts are evidenced with stars (p < 0.05). (B) Mean phase distributions of the group 1 (blue) and group 2 (pink). The two separate populations were sorted based on their individual preferred phase at the second tones (blue and pink dots), where visual entrainment supposedly took place.

The ANOVA on *d’* scores revealed a significant interaction between tone position and group (F(1, 9) = 9.224; p = 0.014) revealing that the visual phase modulation affected differently detection performance between the two groups. Bonferroni-corrected pairwise t-tests showed that this interaction was driven by a greater effect size of tone position in the group 2 than in the group 1, although both groups were better at detecting second tones as compared to first tones. Results also replicated the effect of tone position (F(1, 9) = 12.613; p = 0.006) with greater d’ for the tones sorted as second than first. Finally, no main effect of group was found (F(1, 9) = 0.918; p = 0.363). No difference between threshold (SNR_group1_ = 1.39e10^−3^ ± 2.31e10^−3^; SNR_group2_ = 1.43e10^−3^ ± 3.47e10^−3^; T(1, 24) = −0.66; p = 0.95; two-tailed), nor hit rates (hit_group1_ = 0.761 ± 0.003; hit_group2_ = 0.763 ± 0.004; T(1, 24) = −0.23; p = 0.84; two-tailed) were found in the calibration task, ruling out any hearing difference.

### Stimuli analyses

**Figure S3:**
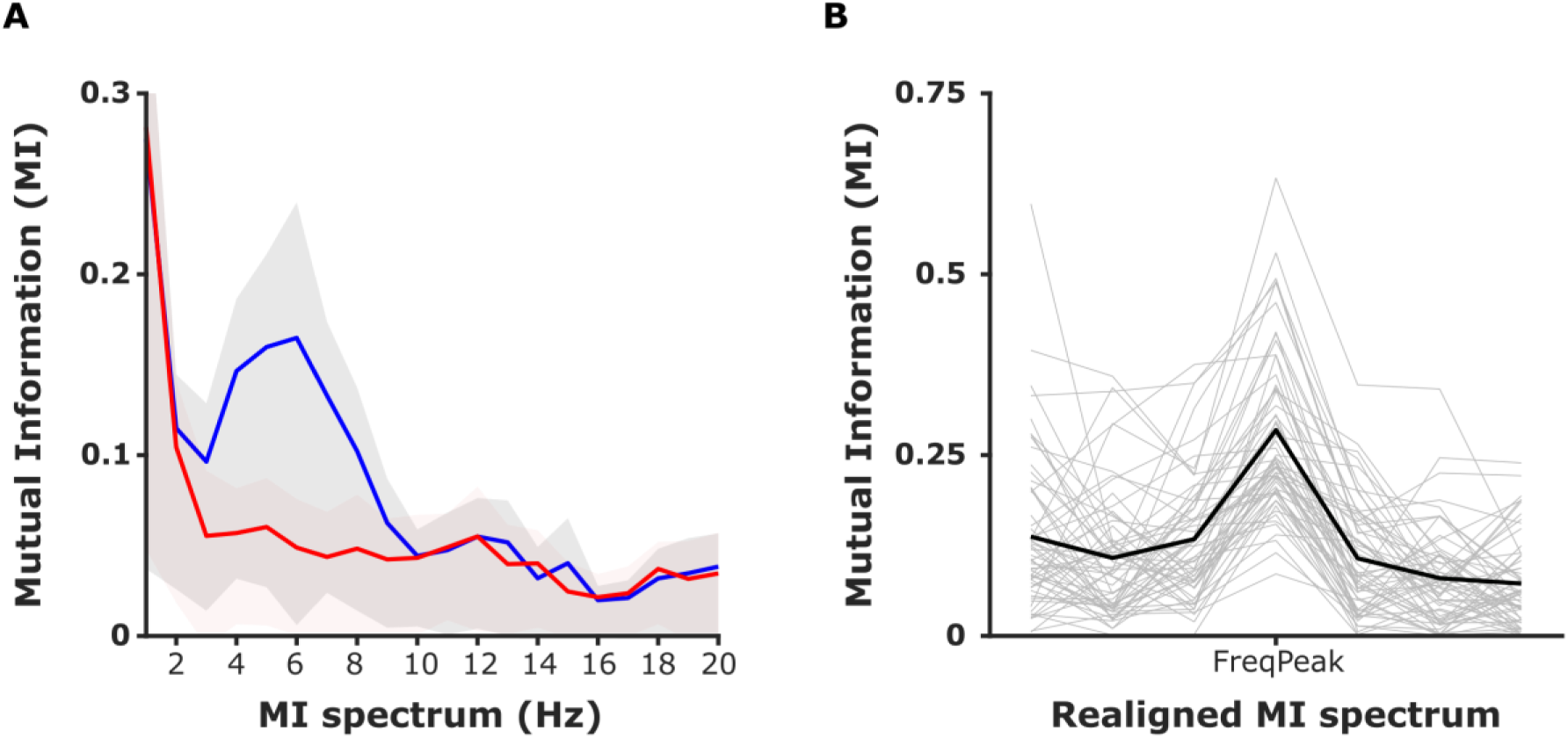
Mutual information between lips movements and auditory envelope in the movies. (A) Mean mutual information spectrum (± standard deviation) between the vertical aperture of the lips and the corresponding (blue line) or random (red line) speech envelope from movies. The greater dependency between the two signals is reflected by the bump localised in the theta frequency band of interest (4 - 8 Hz). (B) Realigned spectrum on the frequency with the greater MI peak (± 1-3 Hz) of each movie (grey lines) and averaged (black line). For each movie, we applied a peak detection and selected the stimuli with a greater MI between the vertical aperture of the lips and auditory envelope situated in the frequencies of interest only (i.e. 4, 5, 6, 7 and 8 Hz).

## REFERENCES

Arnal, L. H., Doelling, K. B., & Poeppel, D. (2015). Delta-Beta Coupled Oscillations Underlie Temporal Prediction Accuracy. Cerebral Cortex, 25(9), 3077–3085. https://doi.org/10.1093/cercor/bhu103

Arnal, L. H., Morillon, B., Kell, C. A., and Giraud, A.-L. (2009). Dual neural routing of visual facilitation in speech processing. J. Neurosci. 29, 13445–13453. doi: 10.1523/jneurosci.3194-09.2009

Assaneo, F., M., & Poeppel, D. (2018). The coupling between auditory and motor cortices is rate-restricted: Evidence for an intrinsic speech-motor rhythm. Sci Adv, 4(2). DOI: 10.1126/sciadv.aao3842

Biau, E., Fernandez, L. M., Holle, H., Avila, C., & Soto-Faraco, S. (2016). Hand gestures as visual prosody: BOLD responses to audio-visual alignment are modulated by the communicative nature of the stimuli. NeuroImage, 132, 129–137

Biau, E., & Kotz, S. A. (2018). Lower beta: A central coordinator of temporal prediction in multimodal speech. Frontiers in Human Neuroscience, Vol. 12. https://doi.org/10.3389/fnhum.2018.00434

Berens, P. (2009). CircStat: A MATLAB Toolbox for Circular Statistics. Journal of Statistical Software, 31(10), 1 – 21. doi: http://dx.doi.org/10.18637/jss.v031.i10

Besle, J., Fischer, C., Bidet-Caulet, A., Lecaignard, F., Bertrand, O., & Giard, M.-H. (2008). Visual activation and audiovisual interactions in the auditory cortex during speech perception: Intracranial recordings in humans. The Journal of Neuroscience, 28(52), 14301–14310. https://doi.org/10.1523/JNEUROSCI.2875-08.2008

Bourguignon, M., Baart, M., Kapnoula, E. C., & Molinaro, N. (2020). Lip-reading enables the brain to synthesize auditory features of unknown silent speech. Journal of Neuroscience, 40(5), 1053–1065. https://doi.org/10.1523/jneurosci.1101-19.2019

Brainard, D.H. (1997). The psychophysics toolbox. Spat. Vis. 10, 433–436

Calvert, G. A., Bullmore, E. T., Brammer, M. J., Campbell, R., Williams, S. C. R., McGuire, P. K., Woodruff, P.W., Iversen, S.D., & David, A. S. (1997). Activation of auditory cortex during silent lipreading. Science, 276(5312), 593–596. https://doi.org/10.1126/science.276.5312.593

Cappe, C., Barone, P. (2005). Heteromodal connections supporting multisensory integration at low levels of cortical processing in the monkey. Eur J Neurosci, 22:2886 –2902

Chandrasekaran, C., Trubanova, A., Stillittano, S., Caplier, A., & Ghazanfar, A. A. (2009). The natural statistics of audiovisual speech. PLoS Computational Biology, 5(7), e1000436. https://doi.org/10.1371/journal.pcbi.1000436

Cogan, G. B., & Poeppel, D. (2011). A mutual information analysis of neural coding of speech by low-frequency MEG phase information. Journal of Neurophysiology, 106(2), 554–563. https://doi.org/10.1152/jn.00075.2011

Crosse, M. J., Butler, J. S., & Lalor, E. C. (2019). Congruent Visual Speech Enhances Cortical Entrainment to Continuous Auditory Speech in Noise-Free Conditions. The Journal of Neuroscience: The Official Journal of the Society for Neuroscience, 35(42), 14195–14204. https://doi.org/10.1523/jneurosci.1829-15.2015

Crosse, M.J, ElShafei, H.A, Foxe, J.J, and Lalor E.C. (2015). Investigating the Temporal Dynamics of Auditory Cortical Activation to Silent Lipreading. 7th Annual International IEEE EMBS Conference on Neural Engineering

Doelling, K. B., Arnal, L. H., Ghitza, O., & Poeppel, D. (2014). Acoustic landmarks drive delta-theta oscillations to enable speech comprehension by facilitating perceptual parsing. NeuroImage, 85 Pt 2, 761–768. https://doi.org/10.1016/j.neuroimage.2013.06.035

Fujioka, T., Ross, B., & Trainor, L. J. (2015). Beta-Band Oscillations Represent Auditory Beat and Its Metrical Hierarchy in Perception and Imagery. Journal of Neuroscience, 35(45), 15187–15198. https://doi.org/10.1523/jneurosci.2397-15.2015

Ghitza, O. (2017). Acoustic-driven delta rhythms as prosodic markers. Language, Cognition and Neuroscience, 32(5), 545–561. https://doi.org/10.1080/23273798.2016.1232419

Giraud, A.-L., & Poeppel, D. (2012). Cortical oscillations and speech processing: emerging computational principles and operations. Nature Neuroscience, 15(4), 511–517. https://doi.org/10.1038/nn.3063

Grant, K. W., Walden, B. E., & Seitz, P. F. (1998). Auditory-visual speech recognition by hearing-impaired subjects: consonant recognition, sentence recognition, and auditory-visual integration. The Journal of the Acoustical Society of America, 103(5 Pt 1), 2677–2690. https://doi.org/10.1121/1.422788

Gross, J., Hoogenboom, N., Thut, G., Schyns, P., Panzeri, S., Belin, P., & Garrod, S. (2013). Speech Rhythms and Multiplexed Oscillatory Sensory Coding in the Human Brain. PLoS Biology, 11(12), e1001752. https://doi.org/10.1371/journal.pbio.1001752

Haegens S, Zion Golumbic E. (2018). Rhythmic facilitation of sensory processing: A critical review. Neurosci Biobehav Rev, 86:150–165. doi: 10.1016/j.neubiorev.2017.12.002

Hanslmayr, S., Axmacher, N., & Inman, C. S. (2019, July 1). Modulating Human Memory via Entrainment of Brain Oscillations. Trends in Neurosciences, Vol. 42, pp. 485–499. https://doi.org/10.1016/j.tins.2019.04.004

Hickok, G., & Poeppel, D. (2007, May). The cortical organization of speech processing. Nature Reviews Neuroscience, Vol. 8, pp. 393–402. https://doi.org/10.1038/nrn2113

Hipp, J. F., Hawellek, D. J., Corbetta, M., Siegel, M., & Engel, A. K. (2012). Large-scale cortical correlation structure of spontaneous oscillatory activity. Nature Neuroscience, 15(6), 884–890. https://doi.org/10.1038/nn.3101

Hishida, R., Hoshino, K., Kudoh, M., Norita, M., Shibuki, K. (2003). Anisotropic functional connections between the auditory cortex and area 18a in rat cerebral slices. Neurosci Res, 46:171–182

Huang Y, Dmochowski JP, Su Y, Datta A, Rorden C, Parra LC (2013) Automated MRI segmentation for individualized modeling of current flow in the human head. J Neural Eng, 10:066004

Ince, R. A. A., Giordano, B. L., Kayser, C., Rousselet, G. A., Gross, J., & Schyns, P. G. (2017). A statistical framework for neuroimaging data analysis based on mutual information estimated via a gaussian copula. Human Brain Mapping, 38(3), 1541–1573. https://doi.org/10.1002/hbm.23471

Keitel, A., Gross, J., & Kayser, C. (2018). Perceptually relevant speech tracking in auditory and motor cortex reflects distinct linguistic features. PLoS Biology, 16(3), e2004473. https://doi.org/10.1371/journal.pbio.2004473

Kleiner, M., Brainard, D., Pelli, D., Ingling, A., Murray, R., and Broussard, C (2007). What’s new in psychtoolbox-3. Perception, 36, 1–16

Kösem, A., & van Wassenhove, V. (2017). Distinct contributions of low-and high-frequency neural oscillations to speech comprehension. Language, Cognition and Neuroscience, 32(5), 536–544

Krahmer, E., & Swerts, M. (2007). The effects of visual beats on prosodic prominence: Acoustic analyses, auditory perception and visual perception. Journal of Memory and Language, 57(3), 396–414

Lakatos, P., Karmos, G., Mehta, A. D., Ulbert, I., & Schroeder, C. E. (2008). Entrainment of neuronal oscillations as a mechanism of attentional selection. Science (New York, N.Y.), 320(5872), 110–113. https://doi.org/10.1126/science.1154735

Leek, M. R. (2001). Adaptive procedures in psychophysical research. Perception and Psychophysics, 63(8), 1279–1292. https://doi.org/10.3758/BF03194543

Luo, H., & Poeppel, D. (2007). Phase patterns of neuronal responses reliably discriminate speech in human auditory cortex. Neuron, 54(6), 1001–1010. https://doi.org/10.1016/j.neuron.2007.06.004

Luo, H., Liu, Z., & Poeppel, D. (2010). Auditory cortex tracks both auditory and visual stimulus dynamics using low-frequency neuronal phase modulation. PLoS Biology, 8(8), 25–26. https://doi.org/10.1371/journal.pbio.1000445

Macmillan, N. A., & Kaplan, H. L. (1985). Detection theory analysis of group data: Estimating sensitivity from average hit and false-alarm rates. Psychological Bulletin, 98, 185–199.

Meyer, L., Sun, Y., & Martin, A. E. (2019). Synchronous, but not entrained: exogenous and endogenous cortical rhythms of speech and language processing. Language, Cognition and Neuroscience, 1–11. https://doi.org/10.1080/23273798.2019.1693050

Michelmann S, Bowman H, Hanslmayr S (2016) The temporal signature of memories: identification of a general mechanism for dynamic memory replay in humans. PLOS Biol, 14:e1002528

Moore, B. C. J., Glasberg, B. R., Varathanathan, A., & Schlittenlacher, J. (2016). A Loudness Model for Time-Varying Sounds Incorporating Binaural Inhibition. Trends in Hearing

Morillon, B., Arnal, L. H., Schroeder, C. E., & Keitel, A. (2019, December 1). Prominence of delta oscillatory rhythms in the motor cortex and their relevance for auditory and speech perception. Neuroscience and Biobehavioral Reviews, Vol. 107, pp. 136–142. https://doi.org/10.1016/j.neubiorev.2019.09.012

Munhall, K. G., Jones, J. A., Callan, D. E., Kuratate, T., & Vatikiotis-Bateson, E. (2004). Visual prosody and speech intelligibility: head movement improves auditory speech perception. Psychological Science, 15(2), 133–137

Obleser, J., & Kayser, C. (2019). Neural Entrainment and Attentional Selection in the Listening Brain. Trends in Cognitive Sciences, Vol. 23, pp. 913–926. https://doi.org/10.1016/j.tics.2019.08.004

Oostenveld R, Fries P, Maris E, Schoffelen JM (2011) FieldTrip: open source software for advanced analysis of MEG, EEG, and invasive electrophysiological data. Comput Intell Neurosci, 2011:156869

Park, H., Ince, R. A. A., Schyns, P. G., Thut, G., & Gross, J. (2015). Frontal Top-Down Signals Increase Coupling of Auditory Low-Frequency Oscillations to Continuous Speech in Human Listeners. Current Biology, 25(12), 1649–1653. https://doi.org/10.1016/j.cub.2015.04.049

Park, H., Ince, R. A. A., Schyns, P. G., Thut, G., & Gross, J. (2018). Representational interactions during audiovisual speech entrainment: Redundancy in left posterior superior temporal gyrus and synergy in left motor cortex. PLoS Biology, 16(8).

Park, H., Kayser, C., Thut, G., & Gross, J. (2016). Lip movements entrain the observers’ low-frequency brain oscillations to facilitate speech intelligibility. ELife, 5. https://doi.org/10.7554/eLife.14521

Peelle, J. E., & Davis, M. H. (2012). Neural Oscillations Carry Speech Rhythm through to Comprehension. Frontiers in Psychology, 3, 320. https://doi.org/10.3389/fpsyg.2012.00320

Peelle, J. E., & Sommers, M. S. (2015). Prediction and constraint in audiovisual speech perception. Cortex; a Journal Devoted to the Study of the Nervous System and Behavior, 68, 169–181. https://doi.org/10.1016/j.cortex.2015.03.006

Pelli, D.G. (1997). The VideoToolbox software for visual psychophysics: transforming numbers into movies. Spat. Vis. 10, 437–442

Pilling, M. (2009). Auditory event-related potentials (ERPs) in audiovisual speech perception. Journal of Speech, Language, and Hearing Research: JSLHR, 52(4), 1073–1081. https://doi.org/10.1044/1092-4388 (2009/07-0276)

Pulvermüller, F., Huss, M., Kherif, F., Del Prado Martin, F. M., Hauk, O., & Shtyrov, Y. (2006). Motor cortex maps articulatory features of speech sounds. Proceedings of the National Academy of Sciences of the United States of America, 103(20), 7865–7870. https://doi.org/10.1073/pnas.0509989103

Rimmele, J. M., Morillon, B., Poeppel, D., & Arnal, L. H. (2018). Proactive Sensing of Periodic and Aperiodic Auditory Patterns. Trends in Cognitive Sciences, 22, 870–882. https://doi.org/10.1016/j.tics.2018.08.003

Shannon C. E. (1948). The mathematical theory of communication. Bell Syst Tech J, 27:379

Smith, Z. M., Delgutte, B., & Oxenham, A. J. (2002). Chimaeric sounds reveal dichotomies in auditory perception. Nature, 416(6876), 87–90. https://doi.org/10.1038/416087a

Tadel F, Baillet S, Mosher JC, Pantazis D, Leahy RM (2011) Brainstorm: a user-friendly application for MEG/EEG analysis. Comput Intell Neurosci, 2011:879716

Thompson, L.,A., Malloy, D. (2004). Attention Resources and Visible Speech Encoding in Older and Younger Adults. Exp Aging Res, 30: 241–252

Thompson, L., A. (1995). Encoding and memory for visible speech and gestures: A comparison between young and older adults. Psychol Aging,10: 215–228

Thorne, J.D., and Debener, S. (2014) Look now and hear what’s coming: on the functional role of cross-modal phase reset. Hear. Res. 307, 144–152

Thut, G., Veniero, D., Romei, V., Miniussi, C., Schyns, P., & Gross, J. (2011). Rhythmic TMS causes local entrainment of natural oscillatory signatures. Current Biology, 21(14), 1176–1185. https://doi.org/10.1016/j.cub.2011.05.049

Tzourio-Mazoyer, N., Landeau, B., Papathanassiou, D., Crivello, F., Etard, O., N. Delcroix, N., Mazoyer, B., & Joliot, M. (2002). “Automated Anatomical Labeling of activations in SPM using a Macroscopic Anatomical Parcellation of the MNI MRI single-subject brain”. NeuroImage, 15 (1): 273–289

Van Veen BD, van Drongelen W, Yuchtman M, Suzuki A (1997) Localization of brain electrical activity via linearly constrained minimum variance spatial filtering. IEEE Trans Biomed Eng, 44:867–880

van Wassenhove, V., Grant, K. W., & Poeppel, D. (2005). Visual speech speeds up the neural processing of auditory speech. Proceedings of the National Academy of Sciences of the United States of America, 102(4), 1181–1186. https://doi.org/10.1073/pnas.0408949102

Wang, D., Clouter, A., Chen, Q., Shapiro, K. L., & Hanslmayr, S. (2018). Single-Trial Phase Entrainment of Theta Oscillations in Sensory Regions Predicts Human Associative Memory Performance. The Journal of Neuroscience: The Official Journal of the Society for Neuroscience, 38(28), 6299–6309. https://doi.org/10.1523/JNEUROSCI.0349-18.2018

Wilson, S. M., Saygin, A. P., Sereno, M. I., & Iacoboni, M. (2004). Listening to speech activates motor areas involved in speech production. Nature Neuroscience, 7(7), 701–702. https://doi.org/10.1038/nn1263

Zoefel, B., Archer-Boyd, A., & Davis, M. H. (2018). Phase Entrainment of Brain Oscillations Causally Modulates Neural Responses to Intelligible Speech. Current Biology, 28(3), 401-408.e5. https://doi.org/10.1016/j.cub.2017.11.071

